# Multidrug resistance plasmids commonly reprogramme expression of metabolic genes in *Escherichia coli*

**DOI:** 10.1101/2023.11.06.565804

**Authors:** Rebecca J. Hall, Ann E. Snaith, Matthew J. N. Thomas, Michael A. Brockhurst, Alan McNally

## Abstract

Multidrug resistant *Escherichia coli* is a leading cause of global mortality. Transfer of plasmids carrying genes encoding beta-lactamases, carbapenamases, and colistin resistance genes between lineages is driving the rising rates of hard to treat nosocomial and community infections. Multidrug resistance (MDR) plasmid acquisition commonly causes transcriptional disruption, and whilst a number of studies have shown strain-specific fitness and transcriptional effects of an MDR plasmid across diverse bacterial lineages, fewer studies have compared impacts of different MDR plasmids in a common bacterial host. As such, our ability to predict which MDR plasmids are the most likely to be maintained and spread in bacterial populations is limited. Here, we introduced eight diverse MDR plasmids encoding resistances against a range of clinically important antibiotics into *E. coli* K-12 MG1655 and measured their fitness costs and transcriptional impacts. The scale of the transcriptional responses varied substantially between plasmids, ranging from >650 to <20 chromosomal genes being differentially expressed. However, neither the scale of regulatory disruption nor the plasmid size correlated with the magnitude of the plasmid fitness cost, which also varied between plasmids. The identities of differentially expressed genes varied among plasmids, although expression of certain metabolic genes and functions were convergently affected by multiple plasmids, including the downregulation of genes involved in L-methionine transport and metabolism. Our data show the complexity of interaction between host genetic background and plasmid genetic background in determining the impact of MDR plasmid acquisition on *E. coli*.

**Importance:** The increase of infections that are resistant to multiple classes of antibiotics, including those isolates that carry carbapenamases, beta-lactamases, and colistin resistance genes, is of global concern. Many of these resistances are spread by conjugative plasmids. Understanding more about how an isolate responds to an incoming plasmid that encodes antibiotic resistance will provide information that could be used to predict the emergence of MDR lineages. Here, the identification of metabolic networks as being particularly sensitive to incoming plasmids suggests possible targets for reducing plasmid transfer.

## Introduction

*Escherichia coli* is a leading cause of nosocomial antibiotic resistant infections globally (1). Multidrug resistant isolates of *E. coli* have been identified worldwide (2; 3; 4; 5; 6; 7), with many resistance genes encoded on plasmids (8; 9). Understanding more about how these plasmids are disseminated therefore has the potential to reduce nosocomial and community spread of drug resistant pathogens.

The *E. coli* species is hugely diverse both phenotypically and genotypically (10; 11), with MDR lineages found predominantly within phylogroups B2 (sequence types ST131 and ST1193), A (ST167, ST410), and F (ST648) (12; 13; 14; 15; 16; 17). Within these lineages are clones that are defined in part by the MDR plasmids that they are known to carry. The H30Rx clone of ST131, for example, is proficient at acquiring multidrug resistance (MDR) plasmids and is differentiated from other ST131 lineages by the presence of an FII-type plasmid encoding the *bla*_CTX-M-15_ *β*-lactamase (17; 18; 12). ST131-H30Rx has also been identified with plasmids encoding *bla*_CTX-M-14_ or *bla*_CTX-M-27_, albeit to a lesser extent (12). The carriage of this particular *bla*_CTX-M-15_-encoding plasmid is also a key feature of the B4/H24RxC clone of ST410 (15).

This raises the possibility that the nature of the plasmid may be influencing acquisition and therefore the emergence and spread of MDR *E. coli* lineages. The dissemination of *bla*_CTX-M-14_ in phylogroups A, B1, and D in Spain has been driven predominantly by K-type plasmids (19), and the spread of the *β*-lactamase *bla*_CMY-2_ in these phylogroups in China has been shown to be via A/C-type plasmids (20). In contrast, *bla*_CMY-2_ spread in Brazil is not influenced entirely by A/C-type plasmids; there, the K-type and B/O-type were the predominant plasmid types (21). The carbapenemase *bla*_NDM-1_ has been identified on plasmid types including L/M (22), FII (23; 24), N (25), A/C (26), and X3 (27). Another carbapenemase, *bla*_OXA_, is known to be encoded on L/M-type (*bla*_OXA-48_ (26; 28; 29), *bla*_OXA-9_ (29)), HI1B-type (*bla*_OXA-1_ (29), and FII-type plasmids (*bla*_OXA-1_, *bla*_OXA-9_ (29)), for example. MDR genes can also co-exist on plasmids, with known examples including the colistin resistance gene *mcr* -1 with *bla*_OXA-48_ (30), *bla*_CTX-M-55_ (5), or *bla*_CTX-M-65_ (31). MDR plasmids are therefore highly variable, and it is important to ask to what extent this influences plasmid dissemination.

There is evidence that the genetic background of the recipient strain plays a role in determining the success of MDR plasmid transfer. This is somewhat evident in the lineage-specific nature of resistant *E. coli* strains, with ST131 clade C (12), ST648 (32; 33; 34), ST1193 (6; 35; 36), and ST410 (37; 38; 15) particularly prone to acquiring resistance. More definitively, recent work by Dunn *et al*introduced an identical *bla*_CTX-M-15_ FII-type plasmid into a diverse set of *E. coli* strains from across the species phylogeny (39). This work showed that the transcriptional response to plasmid acquisition was largely strain-dependent, with even closely related clade C strains exhibiting very different levels of transcriptional response to acquiring the plasmid.

The effect of plasmids on gene expression has also been investigated in other genera. Plasmids have been shown to alter gene expression in *Pseudomonas aeruginosa* PAO1, most notably of metabolic genes (40). In a multidrug resistant strain of *Acinetobacter baumannii*, the presence of a conjugative plasmid resulted in down-regulation of genes involved in biofilm formation and an upregulation of genes related to glutamate/aspartate transport and iron uptake (41). When carrying a plasmid encoding a carbapenemase, carbohydrate metabolism and multidrug efflux systems genes were upregulated in hypervirulent *Klebsiella pneumoniae*, and genes including virulence factors were downregulated (42). These data suggest metabolic networks may be particularly sensitive to altered expression following plasmid acquisition, but whether the nature of the plasmid dictates this when the host background is controlled is not clear.

Here, we determine the extent to which plasmid genetic background can affect the transcriptional response of *E. coli*acquiring MDR plasmids. We use the term ‘MDR’ here as a proxy for the presence of an extended-spectrum beta-lactamase (ESBL), a carbapenemase, or a colistin-resistance gene as these generally co-occur with genes encoding resistance to other antibiotics (43; 44), although we recognise that this is not always the case. We introduced eight different MDR plasmids into *E. coli* K-12 MG1655 and measured the transcriptional response using RNA sequencing. The plasmids represent different replicons, different MDR gene types on similar plasmid types, and plasmids with multiple AMR gene types. We found plasmid specific differential gene expression in response to plasmid acquisition in *E. coli*, and that the level of response did not directly correlate to impacts on fitness. We show that metabolic gene expression is widely affected by MDR plasmid acquisition, with parallelisms in certain pathways and plasmid-specific effects in others, highlighting an important commonality in how MDR plasmids influence transcription in *E. coli*.

## Methods

### Bacterial strains and plasmids

The recipient strain was *E. coli* K-12 MG1655. Eight different plasmid-bearers (*E. coli*, *K. pneumoniae*) were used as donors to transfer different MDR plasmids by conjugation. *Klebsiella pneumoniae* Ecl8 was used as an intermediate when the donor was *E. coli*. Here, we use ‘MDR plasmid’ as proxy for plasmids that carry an ESBL, carbapenemase, or colistin resistance gene. Donor strains and their corresponding MDR plasmids are detailed in Table 1.

**Table 1.**
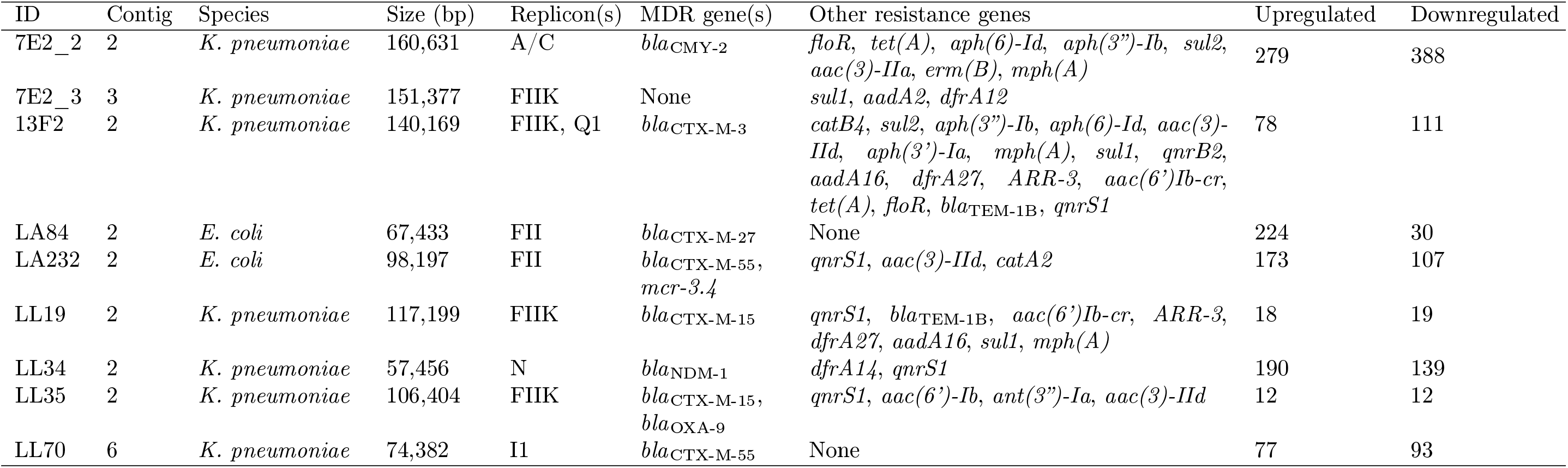
The conjugative plasmids and resulting differential gene expression, including plasmid ID used throughout, the contig on which the plasmid is predicted to be located, the donor species, the predicted size (base pairs [bp]) of the plasmid, the plasmid replicon(s), MDR gene(s), other predicted resistance genes, and the total number of genes upregulated and downregulated in the transconjugants relative to the plasmid-free strain after a false discovery rate threshold of p<0.05 and an absolute log fold change of at least 1 was applied. Resistance genes were considered present and reported here if they were scored by ABRicate as over 80% coverage and over 95% identity. All plasmid replicons reported by ABRicate using default cut-offs are reported here after confirmation using PubMLST.

### Plasmid analysis

Plasmid replicons and plasmid-encoded resistance genes were predicted using ABRicate (v0.8) to query the PlasmidFinder and ResFinder databases, respectively, from the long-read (LA232, LL19, LL35) and hybrid (13F2, 7E2, LA84, LL34, LL70) assemblies. Plasmid types were confirmed using PubMLST (45). This information, alongside plots generated using Bandage (v0.8.1), was used to identify the contigs that contained the plasmids. These were separated from the rest of the assembly using TigSPLIT (https://github.com/stevenjdunn/TigSPLIT) and annotated using Prokka (46) (v1.14.6). Plasmid maps were generated using SnapGene (v6.1.0). A pangenome of the plasmids was constructed using Panaroo (47) (v1.2.10) with default parameters and a sensitive clean mode. A phylogeny of the plasmid sequences was generated using MashTree (v1.2.0), and this plus the gene_presence_absence_roary output from Panaroo was visualised using Phandango (48).

### Assigning function to plasmid-encoded genes

Functions were assigned to genes present in the contigs containing the plasmids using the method described in (49). Briefly, the pangenome reference file produced by Panaroo was translated from nucleotide into peptide sequence using the CD-SToAminoAcids.py custom Python script from https://github.com/C-Connor/GeneralTools. Functional annotation was performed using eggNOG-mapper (v2.1.9) (50) based on eggNOG orthology data (51) to assign COG categories to each gene. Sequence searches were performed using DIAMOND (v2.0.15). Genes that could not be assigned a functional category were designated ‘-’. For visualisation purposes, genes that were assigned a mixed function category (NU, for example) were grouped under ‘Other’.

### Gene Ontology enrichment analysis

To assess which functions were enriched in the differentially expressed (DE) genes, Gene Ontology (GO) enrichment analysis was performed on the significant genes as a whole and for each transconjugant individually using the PANTHER 17.0 classification system, with Fisher’s Exact test and corrected using false discovery rate (52; 53). Hypothetical genes or those without an assigned identification were excluded.

### Plasmid conjugation

Liquid broth conjugations from LL19, LL34, 13F2, and LL35 (as in (54)) donors was performed as per methods detailed in (39). Briefly, an overnight culture was grown in 5 ml LB (*E. coli* MG1655 recipient) or 5 ml LB (E & O Laboratories Ltd) + 4 *µ*g/ml cefotaxime (Alfa Aesar) (donors) at 37*^◦^*C from a single colony. The 5 ml overnight culture was centrifuged at 8,000 rpm (Eppendorf MiniSpin F-45-12-11) for three minutes, the pellet resuspended in 1 ml phosphate buffered saline (PBS, VWR), and this washing step repeated twice before resuspending in a final volume of 500 *µ*l PBS. A sample of 20 *µ*l donor and 80 *µ*l recipient was mixed by brief vortexing before transfer into 5 ml brain-heart infusion (BHI, Sigma) broth. The suspension was inverted to mix and incubated for 18 hours at 37*^◦^*C without agitation. Cultures were then serially diluted in PBS to single colonies and spread on to UTI ChromoSelect chromogenic agar plates (Millipore) containing 4 *µ*g/ml cefotaxime before overnight incubation at 37*^◦^*C. Transconjugants were identified by colour and were then plated on to fresh chromogenic agar plates with 4 *µ*g/ml cefotaxime to check purity. Liquid conjugation from the LA84 donor was performed as described above, with the additional step of conjugation via a *K. pneumoniae* Ecl8 intermediate.

Filter mating conjugations from 7E2, LA232, and LL70 donors was performed by growing an overnight culture in 5 ml LB (*E. coli* MG1655 recipient) or 5 ml LB + 4 *µ*g/ml cefotaxime (donors) at 37*^◦^*C from a single colony. A 1 ml sample of overnight suspension was centrifuged for five minutes at 8,000 rpm (Eppendorf MiniSpin F-45-12-11), the pellet resuspended in 1 ml PBS, and this washing step then repeated twice before resuspending in a final volume of 500 *µ*l PBS. Sterile filter paper discs were placed on to LB agar plates before 10 *µ*l donor and 10 *µ*l recipient added to each. Plates were inverted and incubated for seven hours at 37*^◦^*C. Filter discs were removed with sterile forceps and vortexed thoroughly in 1 ml PBS, serially diluted in PBS to single colonies, and 100 *µ*l spread on to chromogenic agar plates containing 4 *µ*g/ml cefotaxime. Agar plates were incubated overnight at 37*^◦^*C before the presence of pure transconjugants was assessed as for previous conjugations. Filter conjugation from the LA232 donor was performed as described above, with the additional step of conjugation via a *K. pneumoniae* Ecl8 intermediate. All conjugations were performed in biological triplicate.

### Genome sequencing

Long-read sequencing of the transconjugants containing plasmids from the LA232, LL19, and LL35 donors was performed using MinION sequencing (Oxford Nanopore Technologies, UK). Genomic DNA was extracted from overnight cultures using the Monarch Genomic DNA Purification Kit (New England Biolabs). DNA was quantified using a Qubit 4 fluorometer (Invitrogen) and accompanying broad-range double stranded DNA assay kit (Invitrogen). Sequencing libraries were prepared using SQK-LSK109 ligation sequencing kit and EXP-NBD104 native barcode expansion (Oxford Nanopore Technologies, UK), as per manufacturer instructions. Long-read sequencing was performed on a MinION sequencer using an R9 flow cell (Oxford Nanopore Technologies, UK). Sequences were base called using Guppy (v6.0.1). Reads were filtered using Filtlong (v0.2.1) using a cut-off of 600000000 target bases, demultiplexed using qcat (v1.1.0), and assembled using Flye (v2.8.3-b1695). Hybrid assemblies of the transconjugants containing plasmids from the 7E2, 13F2, LA84, LL34, and LL70 donors were generated by MicrobesNG using their enhanced sequencing service. A hybrid assembly for the plasmid-free parent strain has been published previously (55).

### Transcriptome sequencing

RNA sequencing was performed on biological triplicates of the plasmid-free and the transconjugant strains. For sample preparation, a single colony for each replicate was picked following overnight growth on LB agar and added to 5 ml of LB broth (Sigma-Aldrich, United Kingdom). LB was supplemented with 4 *µ*g/ml cefotaxime for transconjugant culture. A 50 *µ*l suspension of each overnight culture was then transferred into 5 ml fresh LB/LB + cefotaxime media and incubated at 37*^◦^*C with agitation until an optical density at 600 nm (OD600) of approximately 0.85. A 1.5 ml sample was centrifuged for five minutes at 8,000 rpm (Eppendorf MiniSpin F-45-12-11), resuspended in 1 ml phosphate buffered saline (PBS), and this wash step repeated. The supernatant was aspirated and the pellet frozen prior to processing and RNA sequencing by GENEWIZ from Azenta Life Sciences (Frankfurt, Germany) using their standard RNA sequencing service. Differential gene expression was quantified using Kallisto (v0.48.0) The long-read assembly, annotated using Prokka (v1.14.6), of the plasmid-free strain was used as a reference. The annotated assembly was processed using genbank_to_kallisto.py (https://github.com/AnnaSyme/genbank_to_kallisto.py). GNU parallel (56) was used for job parallelization. Differential gene expression was analyzed using the Voom/Limma method in Degust (v4.1.1) with a false discovery rate threshold of p*<*0.05 and an absolute log fold change of at least one. EcoCyc (57), Uniprot (58), and KEGG (59) databases were used to identify gene function and associated biochemical pathways.

### Relative fitness assay

We conducted pair-wise competitions to assess the relative fitness of the plasmid bearers compared to GFP-labelled plasmid-free MG1655 (60). Overnight cultures were created from LB agar plates in 10 ml LB (plasmid-free) and 5 LB with 4 *µ*g/ml cefotaxime (transconjugants) from single colonies of the plasmid-free strain and the biological triplicates of the transconjugants. All cultures were diluted to approximately 1 x 10^9^ cells/ml in PBS. Transconjugant cultures were then centrifuged for five minutes at 10,000 rpm (Eppendorf MiniSpin F-45-12-11), resuspended in 1 ml PBS, centrifuged for five minutes at 10,000 rpm, and resuspended in 1 ml LB. Plasmid-free cultures were then centrifuged for ten minutes at 3,600 rpm (Thermo Scientific Megafuge 40R TX-1000), resuspended in 28 ml PBS, centrifuged for ten minutes at 3,600 rpm, and resuspended in 28 ml LB. All cultures were then serially diluted to approximately 1 x 10^5^ cells/ml before 50 *µ*l plasmid-free and 50 *µ*l transconjugant were mixed in 5 ml LB and incubated for 24 hours at 37*^◦^*C. Samples of 0 hour (hr) and 24 hr mixed populations were stored at -80*^◦^*C until flow cytometry analysis.

### Flow cytometry

Plasmid-free and plasmid-bearers pre- and post-competitions were quantified using the Attune NxT flow cytometer and its associated software package. Prior to analysis, 2 ml of the 0 hour (hr) and 1 ml of the 24 hr mixed populations were thawed and washed three times for five minutes at 12,000 rpm in 1 ml sterile, filtered HEPES buffer solution (Gibco). The whole populations were then stained using 10 *µ*M SYTO^TM^84 (Invitrogen) as per (61), thus resulting in single (SYTO^TM^84) stained transconjugants and double (SYTO^TM^84 plus GFP) stained plasmid-free cells. SYTO^TM^84 was visualised using a 561 nm yellow laser and 585/16 nm emission filter (YL1 channel) (61). GFP was visualised with a 488 nm blue laser and 530/30 nm emission filter (BL1 channel). Example gating is given in Fig. S1, and raw .fcs files are available at 10.6084/m9.figshare.24476680. A competition index (CI) was calculated as per the Miles and Misra method (62), whereby a CI larger than one represented competition in favour of the plasmid-free strains. The Pearson correlation coefficient to measure the relationship between total number of DE genes and CI was calculated using pearsonr from the SciPy package.

## Results

### Collating a panel of diverse conjugative plasmids encoding MDR genes

A panel of donors carrying diverse conjugative plasmids was selected. The plasmids are of variable sizes, ranging from 160,631 bp (p7E2_2) to 57,456 bp (pLL34) (Table 1). They encode MDR genes including *bla*_NDM-1_ (pLL34), *bla*_OXA-9_ (pLL35), *mcr-3.4* (pLA232), and various *bla*_CTX-M_ genes, and possess replicons including FII, N, and I (Table 1, Fig. 1). Eight unique transconjugants were generated in biological triplicate using *E. coli* K-12 MG1655 as a recipient. One transconjugant (7E2) was found to have taken up two plasmids, only one of which was classified as an MDR plasmid (p7E2_2) (Fig. 1).

**Fig. 1.**
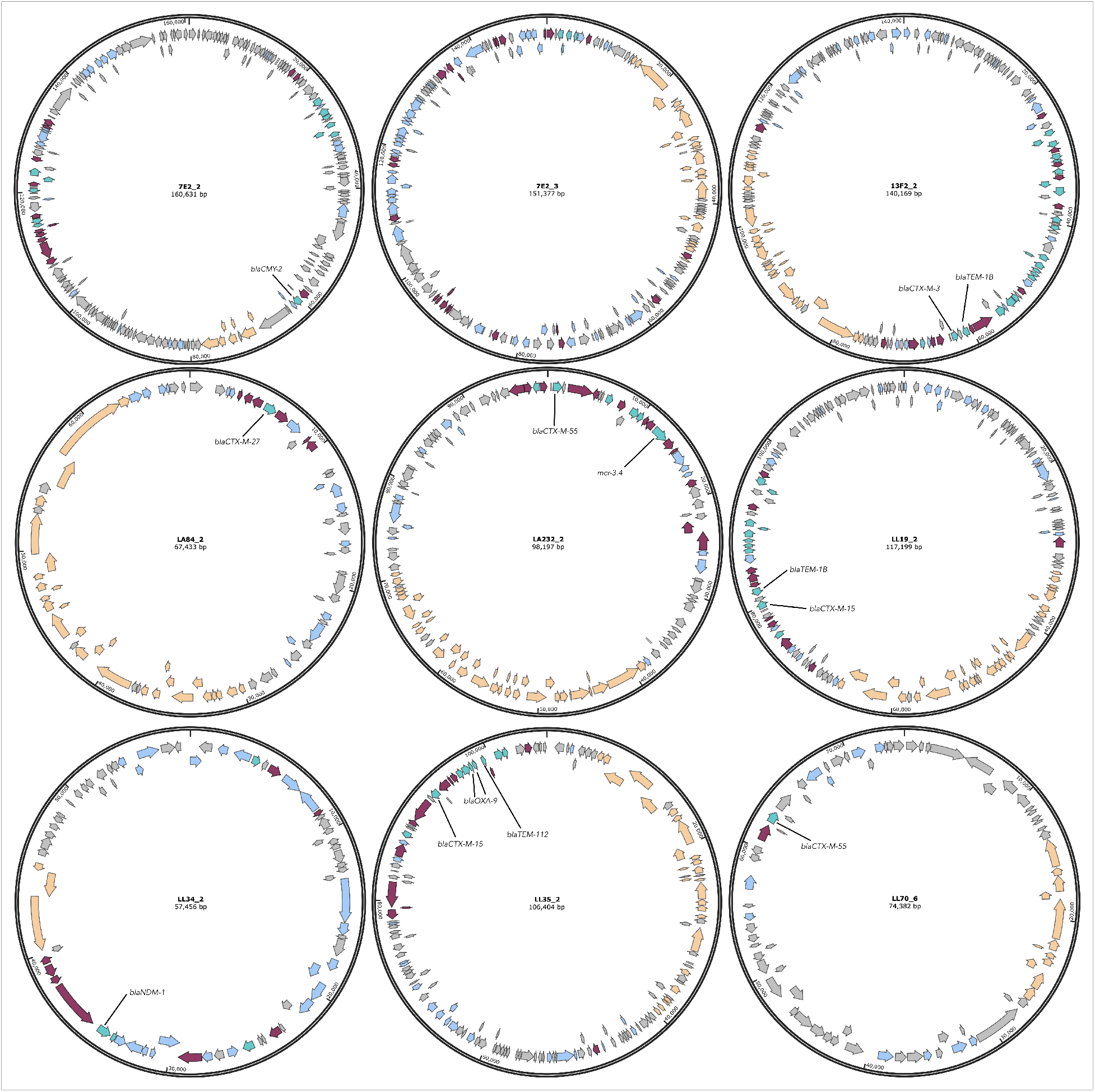
Maps for the nine plasmids conjugated into *E. coli* MG1655. Turquoise = resistance genes, including ones not identified by ResFinder. Orange = conjugal transfer genes, including potential gene/gene fragments not identified by Prokka but found using manual BLAST queries. Maroon = transposase. Grey = hypothetical protein. Blue = annotated gene of other function.

The remaining plasmid-encoded genes are also variable (Fig. S2), with many having unknown function (Fig. 2). Of those to which a COG functional category could be assigned, the largest proportion on each plasmid are involved in replication, recombination and repair (Fig. 2). The only exception to this is the plasmid encoded by the LL70 transconjugant, for which the largest proportion of known genes have functions that straddle several COG categories. Genes involved in inorganic ion transport and metabolism are particularly abundant on p7E2_3 (n=14) in comparison to the other plasmids (n=0 for pLL70, n=1 for pLA232, pLL34, pLL35). Other genes relating to transport and metabolism are scarce on all plasmids. Overall, whilst some plasmids do share some genes, the majority of genes appear to be specific to the individual plasmid (Fig. 2). These plasmids therefore provide a suitably diverse panel from which to examine the immediate transcriptional response upon their acquisition.

**Fig. 2.**
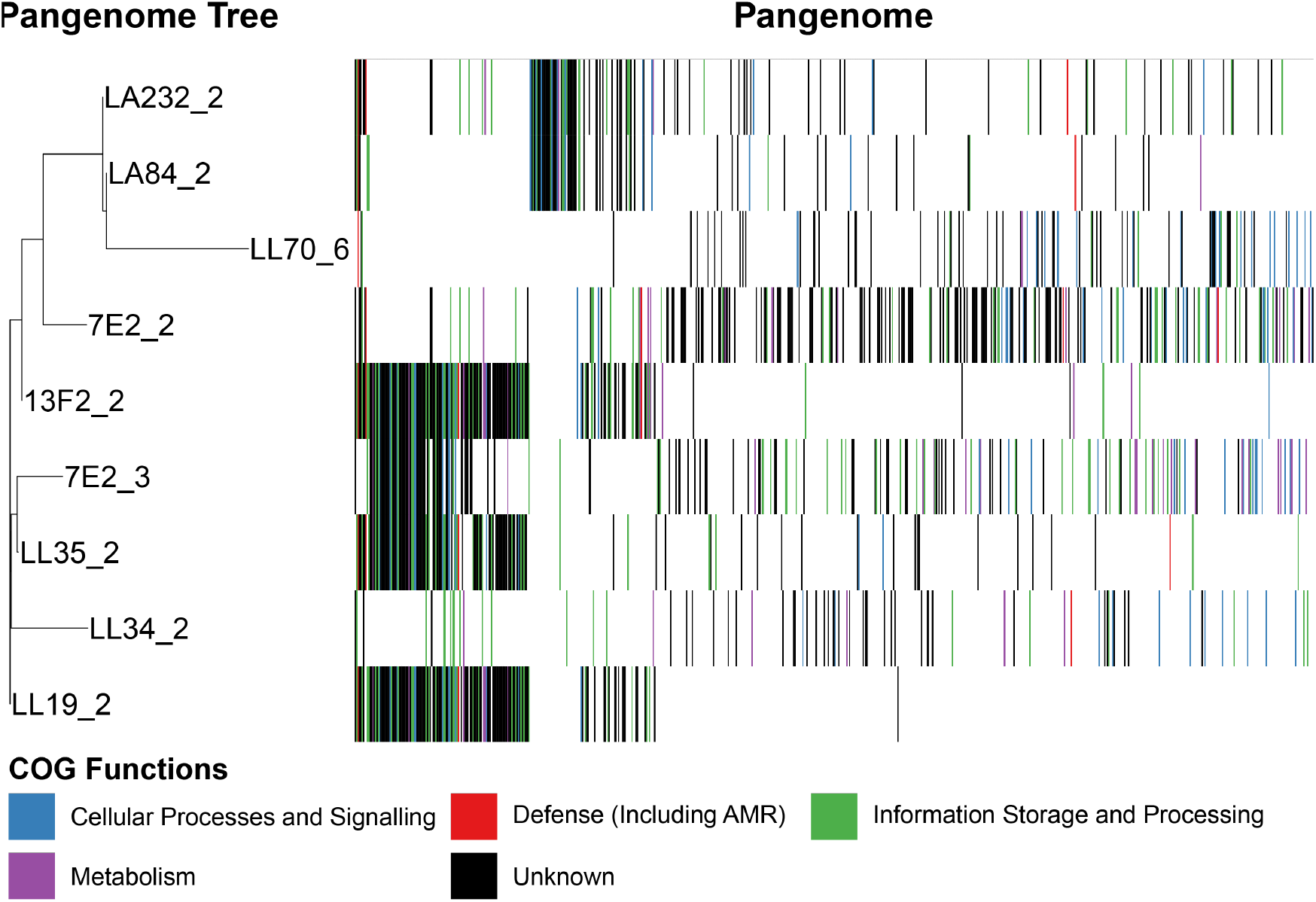
Presence and absence of plasmid-encoded genes on the nine conjugated plasmids. Genes coloured by functions as assigned by eggNOG, grouped into categories for simplicity.

### Scale of transcriptional response does not correlate to plasmid size or fitness cost

Gene expression of the eight different transconjugants was analysed by RNA sequencing to assess the immediate transcriptomic impact of MDR plasmid acquisition. We uncovered large variability in the number of differentially expressed (DE) genes in response to plasmid acquisition (Fig. 3). For example, acquisition of the N-type pLL34 (*bla*_NDM-1_) and the FII-type pLA232 (*bla*_CTX-M-55_, *mcr-3.4*) resulted in relatively high numbers of upregulated genes (190 and 173, respectively), and the largest transcriptional response was observed on acquisition of p7E2, an A/FII plasmid encoding *bla*_CMY-2_, both with regards to upregulated (n=279) and downregulated (n=388) genes. There was no consistent pattern between the transconjugants as to whether there were more upregulated genes than downregulated, or vice versa (Table 1).

**Fig. 3.**
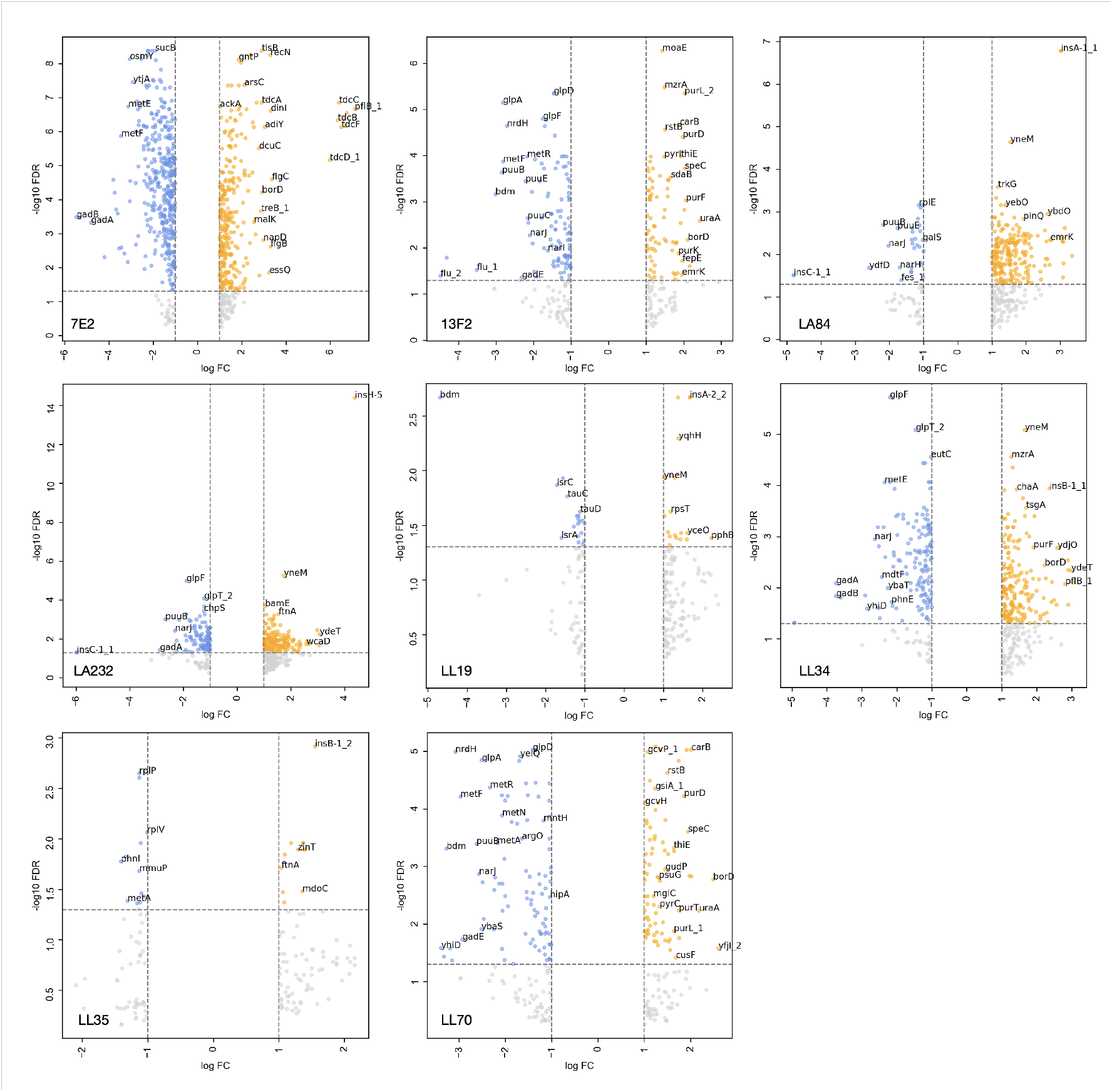
Genes significantly upregulated (orange) and downregulated (blue) (false discovery rate [FDR] threshold of p*<*0.05 and an absolute log fold change [FC] of at least one) in the transconjugants in comparison to the plasmid-free control. Genes which had an absolute log FC of at least one but did not reach the FDR threshold are shown in grey and are considered to not be significantly differentially expressed. Plots are labelled by donor with respect to Table 1. Select genes are labelled.

Plasmid size did not affect the scale of differential gene expression. The pLL34 plasmid was the smallest but triggered one of highest number of DE genes (n=329), whereas LA232 also has a high number of DE genes (n=280) after acquiring a much larger plasmid (Table 1). The transconjugant that acquired two plasmids (160,631 bp and 151,377 bp), 7E2, did however show substantially more DE genes (n=667). Additionally, no significant correlation was observed between the fitness cost of the plasmid and the total number of DE genes (Fig. 4, Pearson correlation coefficient, p = 0.62). Together, these data suggest that transcriptional response to MDR plasmid acquisition in MG1655 is likely influenced by multiple factors, including the presence of other plasmid-encoded gene, rather than one primary driver.

**Fig. 4.**
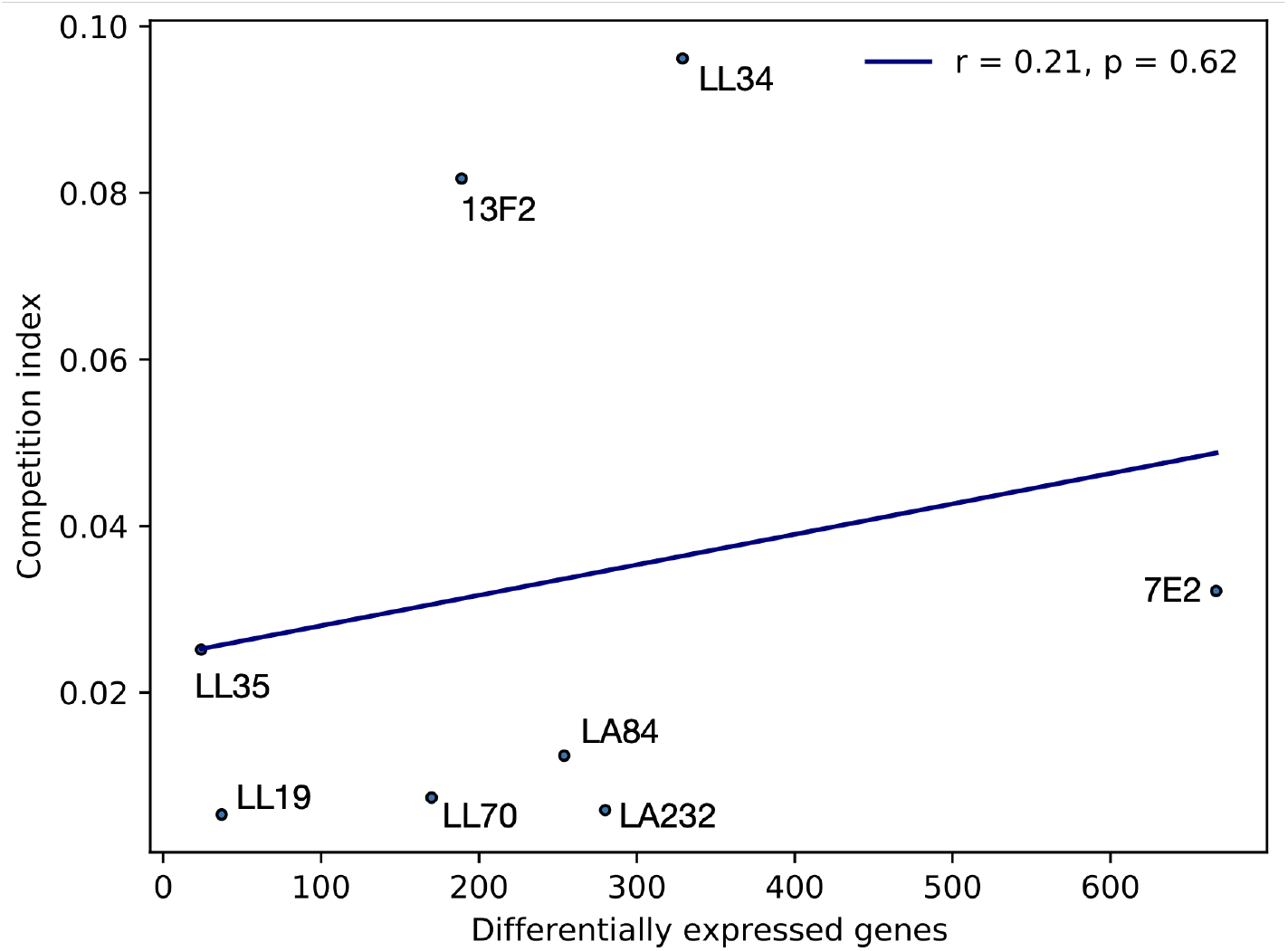
Correlation (Pearson) between total number of differentially expressed genes and competition index of the transconjugants against a plasmid free strain (mean of biological triplicate), whereby a value greater than one indicates the transconjugant is fitter in the competition condition.

### Majority of differentially expressed genes are plasmid specific

We then analysed the significantly DE genes to identify commonalities across the transconjugants. Only one gene (*rpsT*, encoding the 30S ribosomal subunit protein S20) was significantly upregulated in all eight transconjugants (Fig. S3). 34 genes were upregulated in parallel in LA84, LA232, and LL34, including *argACF* (L-arginine biosynthesis from L-glutamate via L-ornithine), *holE* (DNA polymerase III subunit), and *ydiM* (inner membrane transport protein). 21 genes were significantly DE in both LA84 and LA232, including *crcB* (fluoride-specific ion channel, *citC* (citrate lyase synthetase), and genes encoding uncharacterised proteins (*ybbD*, *ybcK*). The number of genes upregulated in one transconjugant exclusively varied between the recipients (7E2, n=131; 13F2, n=3; LA84, n=59; LA232, n=11; LL19, n=3; LL34, n=17; LL35, n=0; LL70, n=16). 17 transconjugant pairs or groups had only one significantly upregulated gene in common, and there were no genes that were significantly downregulated in all eight of the transconjugants. Together, these data suggest that transcriptional response to MDR plasmid acquisition is specific to the incoming plasmid, and that the response to certain plasmids (p7E2) is larger and more plasmid-specific than for others (pLL35).

### Differentially expressed genes enriched in metabolic and transport functions

We performed Gene Ontology (GO) enrichment analysis on the collection of all DE genes across the set of transconjugants to establish an overall pattern. Of the 23 significantly enriched GO terms, nine were related directly to transport (including of metal ions) and a further three to the membrane (Fig. 5, Table S1). Additionally, three GO terms related metabolic processes, covering a total of 136 DE genes across the dataset. This suggests that metabolic processes, including transport, are significantly impacted by the acquisition of MDR plasmids.

**Fig. 5.**
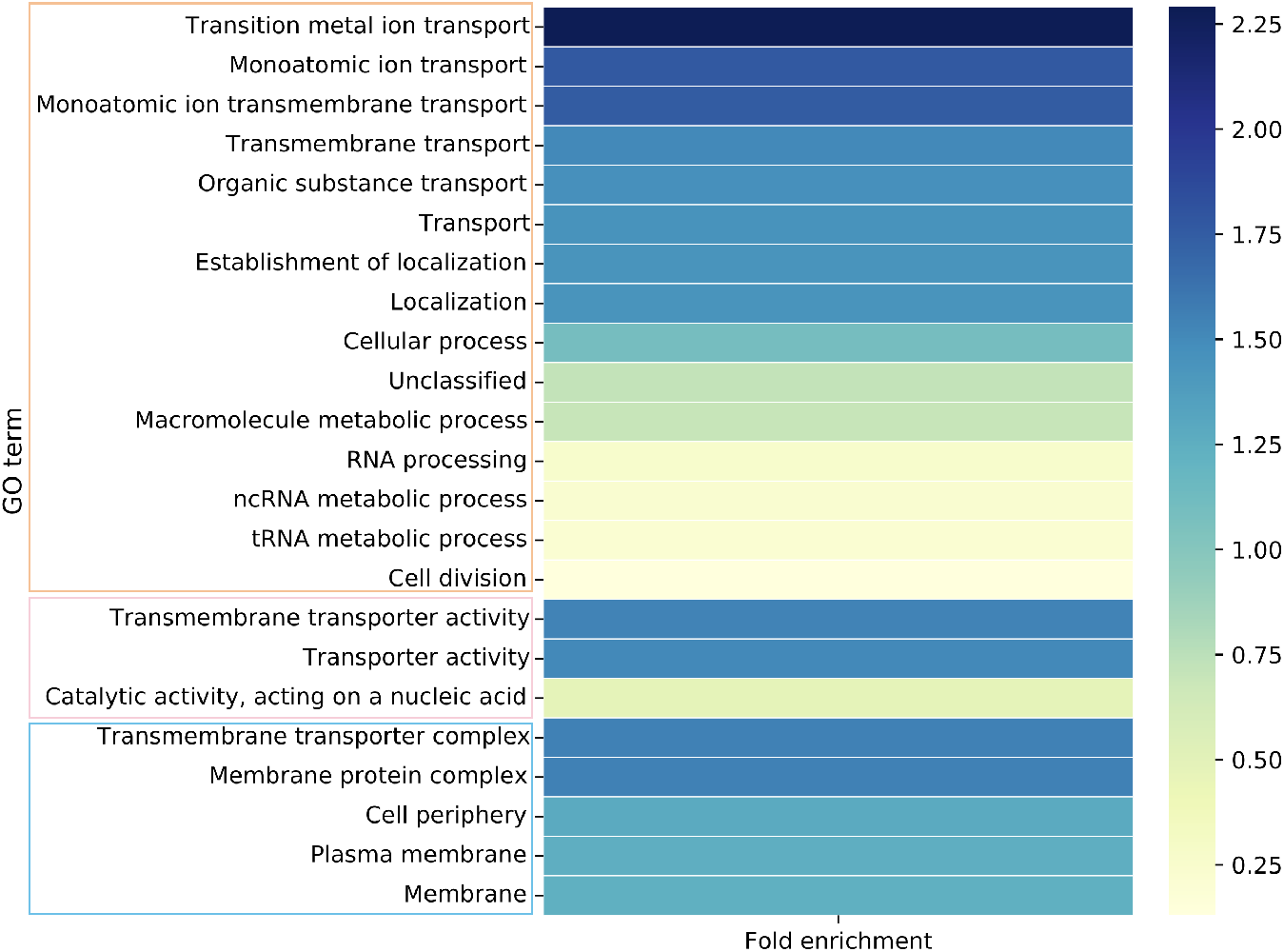
GO terms significantly enriched in the combined set of all transconjugants. A value greater than one indicates more genes observed than expected. Terms are grouped by their role in biological (orange box), molecular (pink), or cellular (blue) function. GO IDs and P values given in Table S1.

### Convergence observed in the downregulation of L-methionine transport and metabolism genes

GO enrichment analysis of the whole transconjugant set identified transport and metabolism as key functions of DE genes. When the GO terms of the DE genes were analysed for each transconjugant individually, five of the eight transconjugants showed enrichment in amino acid metabolism (Supplementary Data 1). This includes aspartate (7E2), methionine (13F2, LL70), arginine (LA232, LL34), glutamate (LL34), and glutamine (LL34, LL70) metabolic pathways, and well as amino acid biosynthesis pathways more generally (7E2, 13F2, LL34, LL70).

The specific patterns of up- or downregulation of genes in these pathways were then examined individually. The 13F2 and LL70 transconjugants showed enrichment in genes involved in L-methionine biosynthetic (13F2, LL70) and metabolic (13F2) processes. Indeed, widespread downregulation in L-methionine transport and metabolism was observed across the transconjugants. The *metA* and *metB* genes that convert L-homoserine to L-cystathionine via O-succinyl-L-homoserine in the production of L-methionine from L-aspartate were downregulated in six transconjugants (Fig. 6). The ATP binding subunit of the MetNI ABC transporter was also significantly downregulated in five transconjugants, whereas the membrane subunit of the same transporter was only downregulated in LL34 (encoding *bla*_NDM-1_). The LA84 transconjugant (*bla*_CTX-M-27_), had no significantly DE genes in this pathway, and only *metB_1* is downregulated in LL19 (*bla*_CTX-M-15_). The convergence observed across this pathway provides evidence towards a hypothesis that downregulation of L-methionine uptake and metabolism may be important in MDR plasmid acquisition and stable integration.

**Fig. 6.**
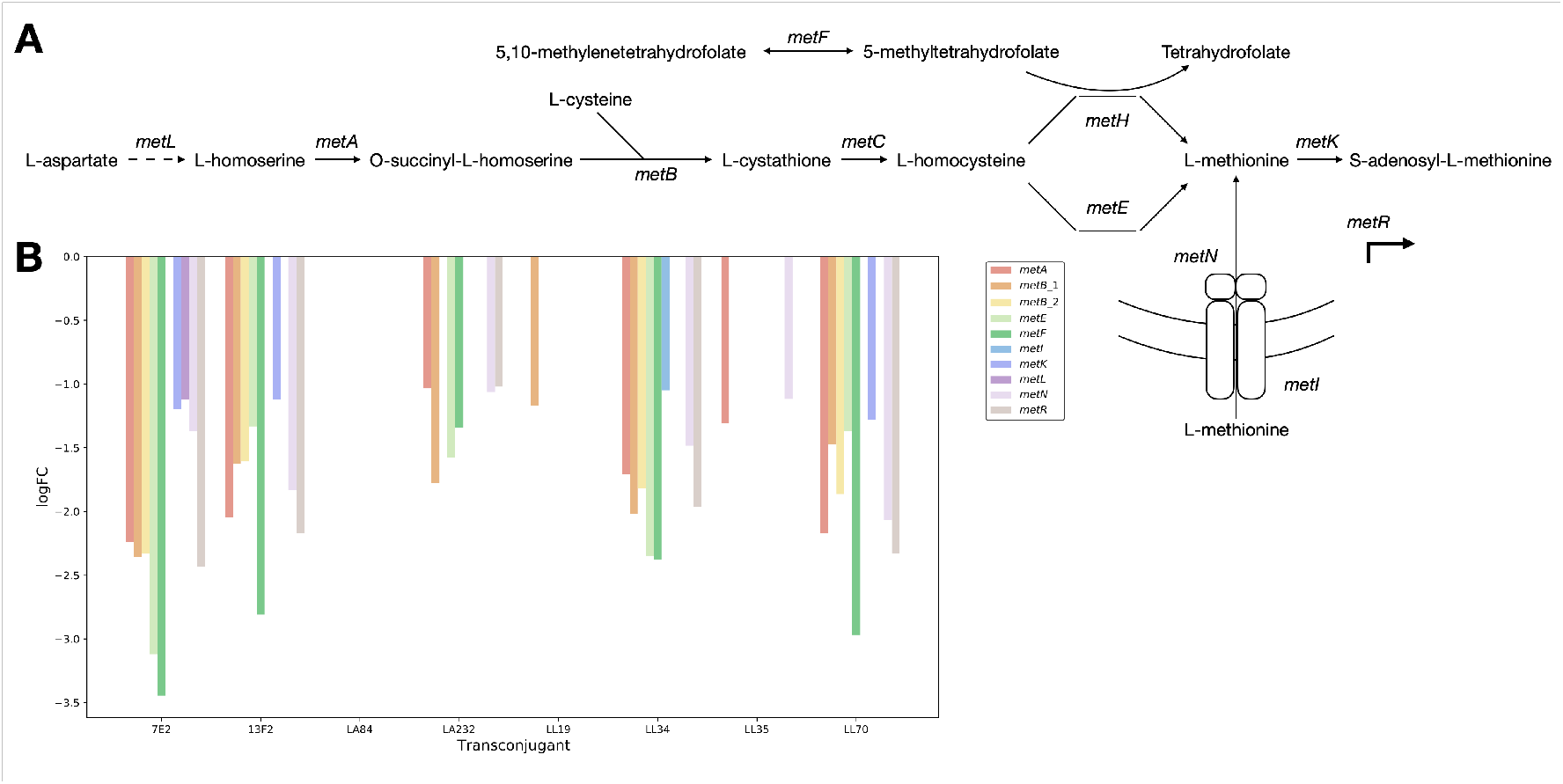
**A** A simplified L-methionine biosynthetic pathway, with key genes highlighted, and **B** the logFC values for the genes downregulated on acquisition of an MDR plasmid, coloured by gene involved in L-methionine transporter and biosynthesis.

### A proposed requirement for L-arginine following plasmid acquisition

GO analysis further highlighted enrichment in L-arginine and L-glutamate metabolic pathways, specifically in LA232 (arginine catabolic processes to glutamate and succinate, arginine metabolic process) and LL34 (arginine and glutamate metabolic processes).

The *argACFI* genes were significantly upregulated in the LA232, LA84, and the LL34 transconjugants (Fig. 7). The *argA* gene encodes N-acetylglutamate synthase that converts L-glutamate into N-acetyl-L-glutamate in the ornithine biosynthesis pathway for the eventual production of L-ornithine from L-glutamate. The *argC* gene (N-acetylglutamylphosphate reductase) functions in the same pathway, and *argF* and *argI* (ornithine carbamoyltransferase chains F and I) catalyse the first step in the conversion of L-ornithine to L-arginine. The periplasmic binding protein of the L-arginine ABC transporter *artJ* is also upregulated in these three transconjugants.

**Fig. 7.**
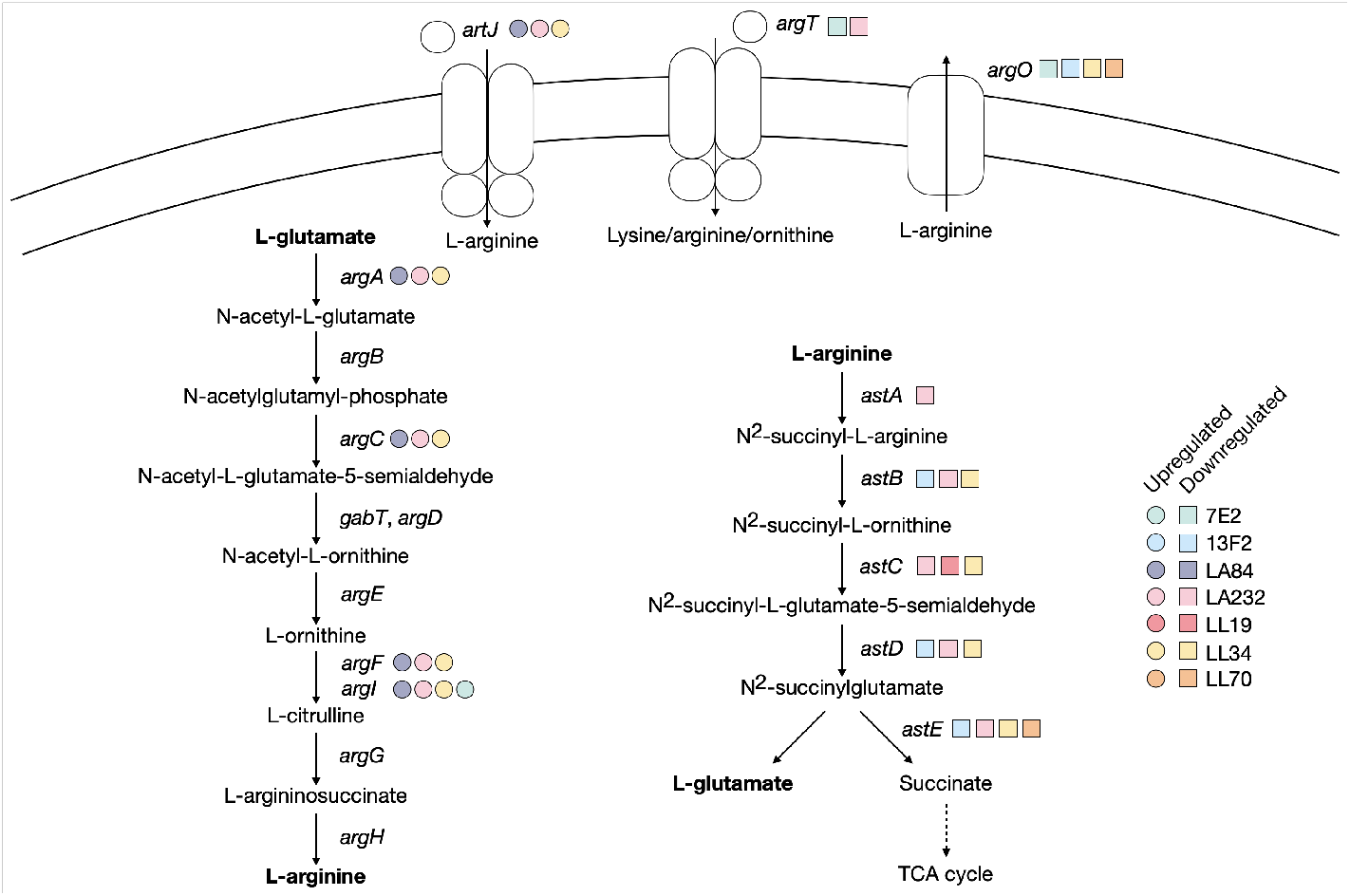
Genes upregulated (circle) and downregulated (square) across all transconjugants that relate to L-arginine biosynthetic pathways. Selected metabolites in each reaction are shown for simplicity, with key metabolites or reactions highlighted in bold.

In contrast, *astABCDE* in the arginine succinyltransferase pathway were downregulated after acquisition of pLA232, and all but *astA* were downregulated in LL34. These genes encode the five enzymes in the pathway that converts L-arginine into L-glutamate and succinate for the TCA cycle. The periplasmic binding protein of the L-ornithine/L-arginine/L-lysine ABC transporter *argT* was also downregulated in LA232 and 7E2, and the L-arginine exporter *argO* was downregulated in LL34, 13F2, 7E2, and LL70. This pattern is not observed in, for example, 7E2, where *argI* is the only common upregulated gene, or in LL35 where none are differently expressed. This suggests two points. First, that the exact metabolic requirement following MDR plasmid acquisition is plasmid-specific. Second, that in response to some MDR plasmids (pLA232, pLL34), there may be a preferential need for L-arginine or a diversion of resources away from the TCA cycle.

### Plasmid-specific enrichment in ATP-binding cassette transporters

Further GO enrichment analysis identified significant differential expression in genes relating to transport across five of the eight transconjugants (with the exception of LL34 and LL35). The nature of the transporters, both in terms of specificity and type, vary between the transconjugants with some evidence of commonalities.

7E2 and 13F2, for example, show significant differential expression of inorganic ion and cation transmembrane transport, with changes also in transition metal and iron transport in the former (Supplementary Data Sheet). GO terms relating to ATP-binding cassette (ABC) transporters are also enriched in 7E2 and LL19. The majority of these genes are downregulated, including the entire *dppABCDF* and *ugpABCE* transporters in 7E2 (responsible for the transport of peptides and glycerol 3-/2-phosphate, respectively) (Fig. 8). Components of the lysine/arginine/ornithine, ferrice enterobactin, and iron (III) hydroxamate, ABC transporters, amongst others, are also downregulated in 7E2, and *lsrACD* are downregulated in both 7E2 and LL19 upon plasmid acquisition. There were fewer examples of ABC transporter genes being upregulated, but it was noted in components of the maltose, nickel, and D-galactose/methyl-galactoside systems.

**Fig. 8.**
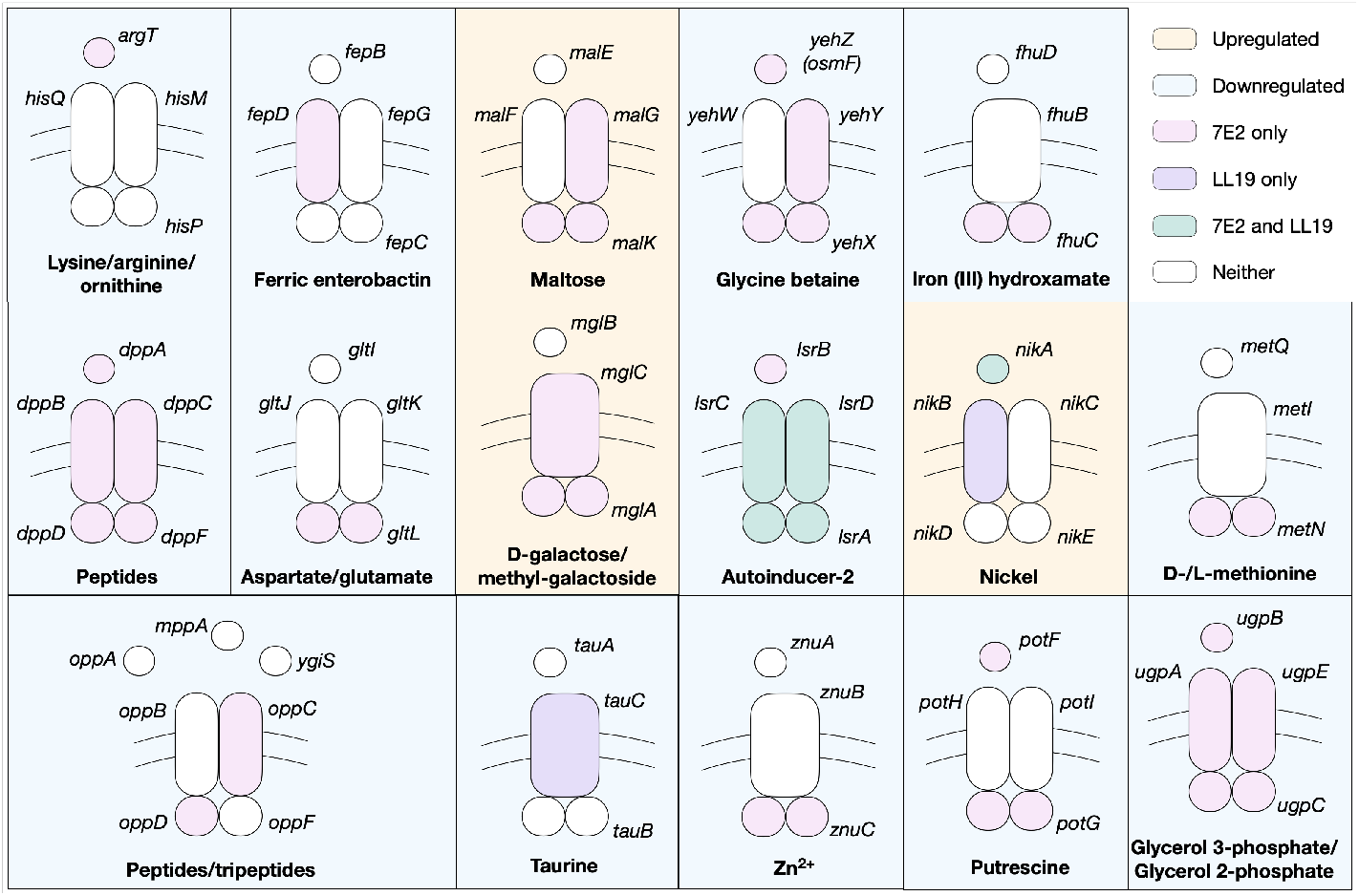
Components of ABC transporters that are significantly upregulated (orange background) or downregulated (blue background) in 7E2 (pink), LL19 (purple), both (green), or neither (white). Simplified schematics of the periplasmic binding, membrane, and ATP-binding proteins are shown. Transporters with significantly DE genes with unknown or putative specificity are not included but are detailed in the Supplementary Data Sheet.

A point of note is the significant downregulation of *potF* and *potG*, encoding components of the putrescine ABC transporter, in 7E2, whereas *plaP*, encoding a putrescine/H+ symporter, is upregulated. The *puuABC* genes from the putrescine degradation pathway are also downregulated (Supplementary Data Sheet). The low affinity of PlaP coupled with evidence that it is has functions relating to type 1 pili (63), suggest that putrescine itself is not a desired metabolite under these conditions.

Together, this suggests a diversion of resources away from high affinity transporters in a plasmid-dependent manner as the cell responds to the incoming plasmid.

## Discussion

The spread of MDR is of increasing concern, particularly within clinical settings. Whilst the mechanisms by which MDR plasmids spread, are well established, less is understood about the role that the recipient plays in plasmid acquisition; specifically, whether the transcriptional response is host or plasmid specific. To begin to unravel this, we introduced eight plasmids into a laboratory strain of *E. coli* via conjugation and measured the resulting transcriptional response. These plasmids encoded a variety of MDR genes, including different *bla*_CTX-M_ genes, and were of different plasmid types.

We uncovered extensive plasmid-dependent differences in transcriptional response to MDR plasmid acquisition, both in terms of absolute number of genes and the nature of the genes themselves. Genes relating to transport and metabolic processes were particularly DE across the transconjugants. Our work supports existing data from other species; in *P. aeruginosa*, plasmids have been shown to trigger differential expression of metabolic genes (40), indicating metabolic networks are particularly sensitive to the effects of incoming plasmids across genera.

We identified transconjugant-specific upregulation of genes involved in the conversion of L-glutamate to L-arginine alongside the downregulation of genes responsible for converting L-arginine into succinate, mostly notably in LA232 although with some specific gene examples in other transconjugants. This suggests that L-arginine might be important immediately following acquisition of certain MDR plasmids. A link between L-arginine and MDR plasmids has been reported previously; specifically, strains carrying an MDR plasmid were serially passaged in the presence and absence of antibiotic selection and the *artP* gene encoding the L-arginine ABC transporter ATP binding protein was found to be the mostly highly mutated gene (54). Whilst we did not find this particular gene to be differentially regulated to the same extent, we did find significant upregulation of *artJ*, encoding the periplasmic binding protein of the same transporter. There is existing evidence of amino acid transport and metabolism genes being consistently upregulated in *A. baumannii*in the presence of antibiotics (64), so an interesting extension of this work may be to measure gene expression immediately following plasmid acquisition in the presence versus absence of antibiotics.

The differential regulation of the pathways relating to L-arginine biosynthesis was particularly notable in the LA232 transconjugant, but the downregulation of components of the L-methionine biosynthesis pathway was common to all but LA84. Whilst current understanding of the link between specific metabolic pathways and MDR is not complete, there are examples of other plasmids affecting the expression of genes relating to L-methionine biosynthesis in *E. coli*. Specifically, the *E. coli* plasmid pEX18Gm has been shown to increase L-methionine biosynthesis via the upregulation of genes including *metH* (20). MetH is involved in the cobalamin-dependent conversion of L-homocysteine, and whilst we did not find significant differential expression of *metH* we did identify a downregulation of *metE*, encoding the cobalamin-independent reaction. This suggests that the presence or absence of methionine may be important immediately following plasmid acquisition in a plasmid and/or strain-specific manner.

Amongst the significantly DE genes, ABC transporters were also notably enriched in two transconjugants (7E2, LL19), suggesting their importance may be plasmid specific. The majority of these genes were down- rather than upregulated, with the implication that metabolite import may not be a priority after plasmid acquisition. Components of the putrescine ABC transporter were downregulated in 7E2. Putrescine is the major polyamine in *E. coli* (65), and polyamines are known to decrease the permeability of the outer membrane in *E. coli* under stress conditions (66). It is possible that the genes were differentially related as part of a stress response or, when considered alongside the downregulation of genes in the putrescine degradation pathway, a diversion of resources away from its use in metabolic pathways. Whilst there is not enough information here to establish the underlying cause, it presents an interesting avenue for future research.

It is already known that plasmids can manipulate host gene expression by plasmid-encoded transcriptional regulators. A plasmid transcriptional regulator has been shown to have wide-reaching effects on a strain of *Pseudomonas fluorescens* via extensive remodelling of the host proteome, including influencing metabolite uptake (67), and other plasmids have been shown to repress expression of the type VI secretion system to aid successful conjugation (41; 68). This manipulation of host behaviour is hypothesised to be an adaptation of the plasmid to increase its own fitness and likelihood of transmission, either by enhancing host competitiveness in a given environment (vertical transmission) or by promoting conjugation (horizontal transmission) (69). Viewed from this perspective, it is possible that the significant DE of metabolic genes across the transconjugants might play a role in increasing the competitiveness of the host outside of a laboratory environment.

A point of note was the relatively small transcriptional response observed in the two transconjugants that acquired the plasmids encoding *bla*_CTX-M-15_ (LL19, LL35). This was notably distinct to those encoding alternative *bla*_CTX-M_ genes. The multidrug resistant lineage of *E. coli* ST131 that harbours *bla*_CTX-M-15_ is arguably the most successful clone of this species in terms of global dissemination (12; 70). This gene is also found in *E. coli* lineages including ST410 (15; 38) and ST1193 (71; 72). We could therefore speculate that the prevalence of plasmid-encoded *bla*_CTX-M-15_ could potentially be due to the small transcriptional response that its acquisition triggers in the recipient.

We now know that are there are strain- and plasmid-specific responses to the acquisition of MDR plasmids. We have shown here and in two previous studies that differential expression of genes involved in metabolism is common during MDR plasmid acquisition (39; 54), but that the exact metabolic pathways affected and to what extent is plasmid-dependent. Recent studies have also shown the importance of metabolic genes in the evolution of AMR in clinical *E. coli* (73). This study therefore contributes to an understanding of the intrinsic link between metabolism and MDR, and should be explored further in existing and emerging pandemic lineages.

## Acknowledgements

Enhanced genome sequencing was provided by MicrobesNG (http://www.microbesng.com). Standard RNA sequencing was performed by GENEWIZ from Azenta Life Sciences. GFP-labelled *E. coli* MG1655 was provided by Michael J Bottery. The Flow Cytometry Platform at the University of Birmingham provided support for flow cytometry experiments. RJH was supported by a University of Birmingham College of Medical and Dental Sciences Research Development Fund award, and by a NERC grant (NE/T01301X/1) awarded to AM.

## Competing interests

The authors declare no competing financial interests in relation to the work described.

## Data availability statement

The DNA and RNA datasets generated and analysed during the current study are available from NCBI BioProject with accession PRJNA1034827.

## Author contributions

RJH: methodology, formal analysis, investigation, data curation, writing - original draft, visualization. AES: investigation, data curation. MT: visualization. MAB: conceptualization. AM: conceptualization, methodology, supervision, funding acquisition. All authors: writing - review & editing.

## Supplementary

**Fig. S1.**
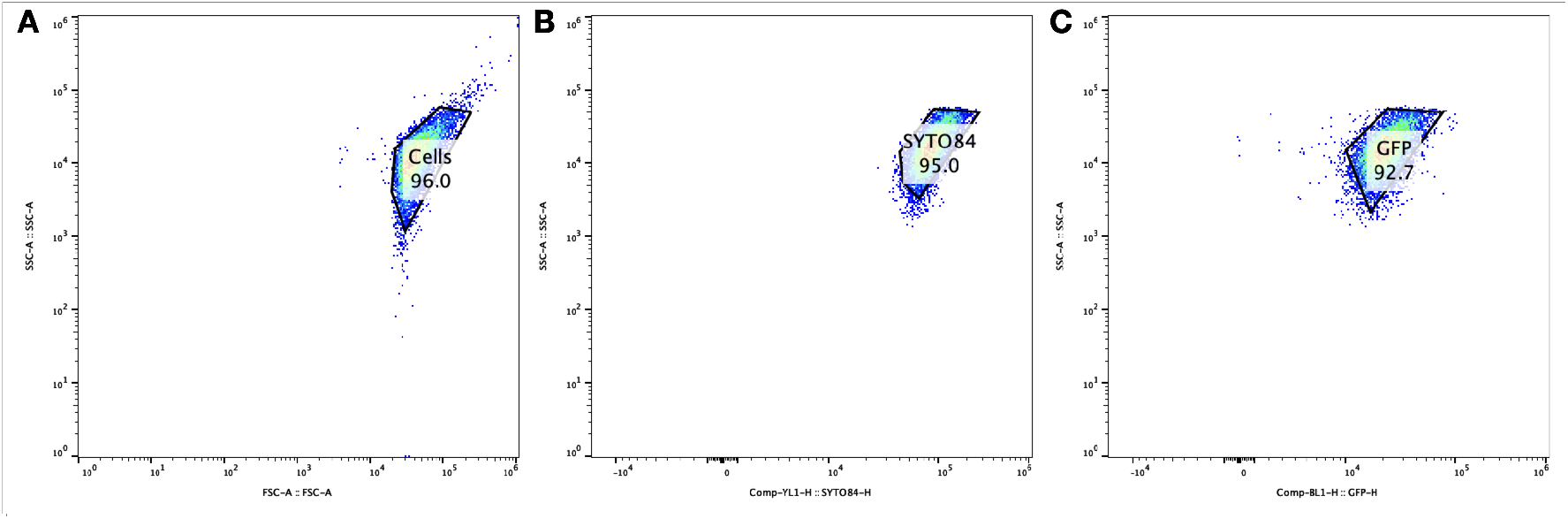
Example gating applied to all test populations. **A** All cells, WT population unstained. **B** SYTO^TM^84 stained cells. **C** GFP+ cells. Gating applied so that single (SYTO^TM^84 only, plasmid bearer) and double (SYTO^TM^84 stained plus GFP+ expressing, plasmid free) labelled cells could be counted.

**Fig. S2.**
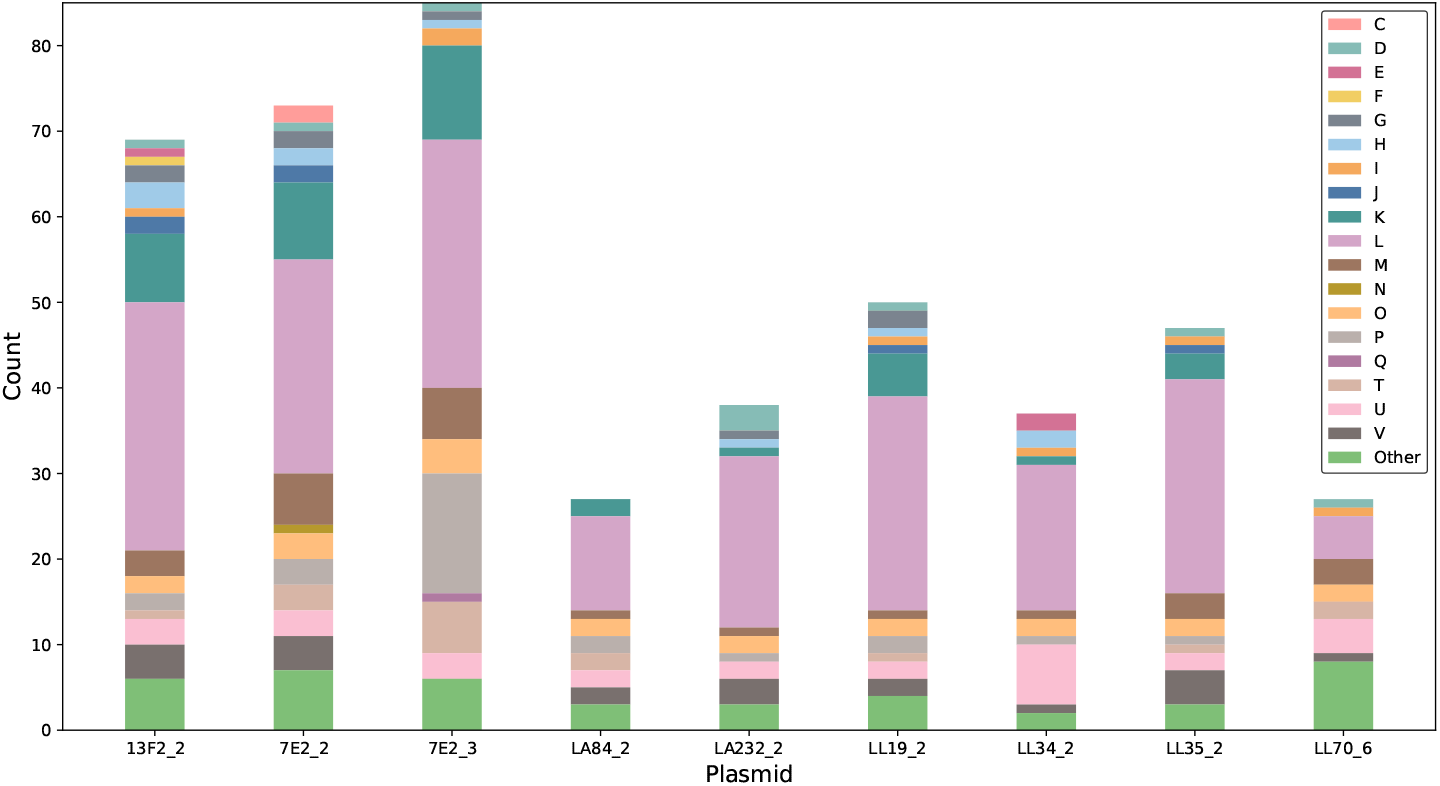
COG categories of genes with known function encoded by the conjugative plasmids, as assigned by eggNOG-mapper.

**Table S1.**
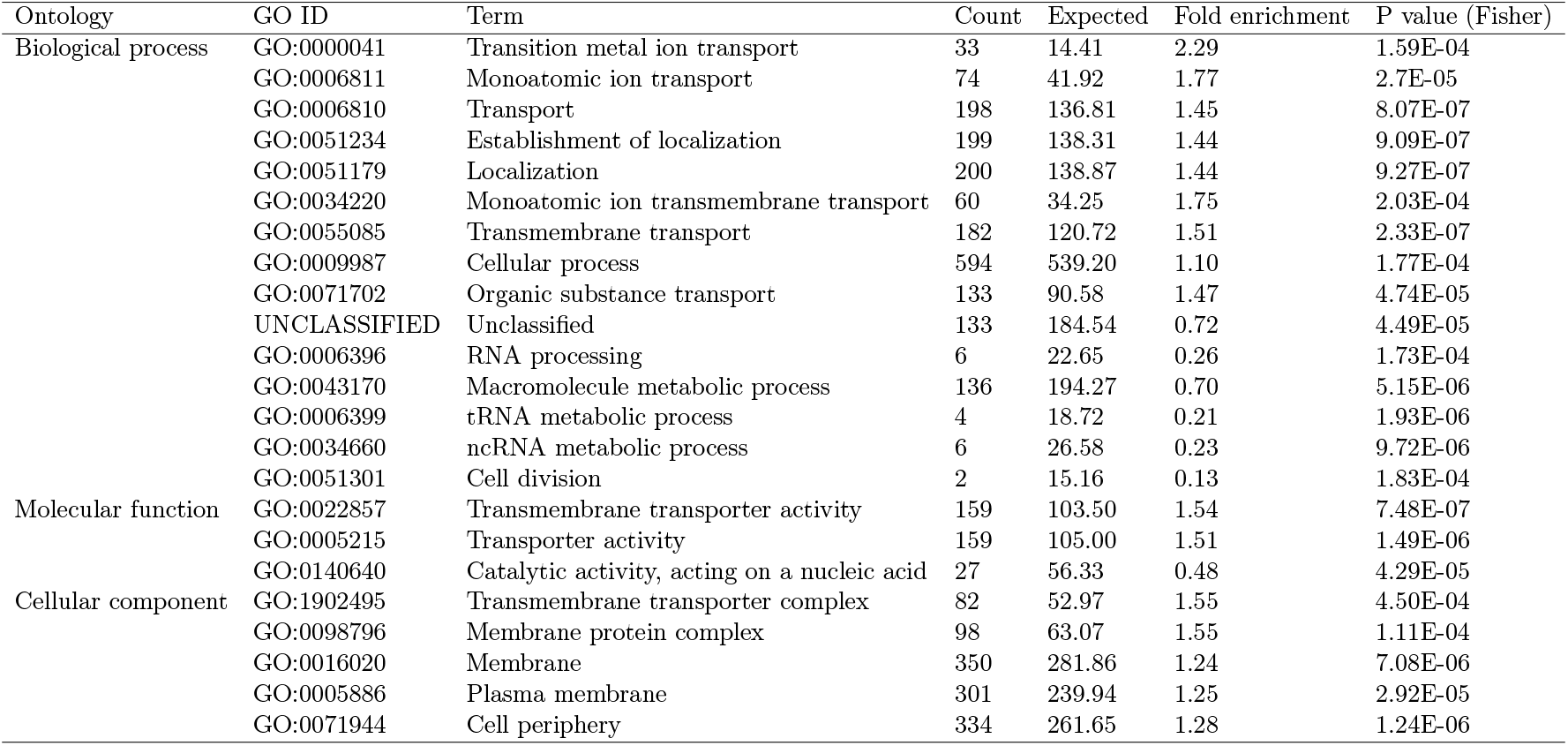
Gene Ontology (GO) terms enriched across the differentially expressed from all transconjugants combined.

| Ontology | GO ID | Term | Count | Expected | Fold enrichment | P value (Fisher) |
| --- | --- | --- | --- | --- | --- | --- |
| Biological process | GO:0000041 | Transition metal ion transport | 33 | 14.41 | 2.29 | 1.59E-04 |
|  | GO:0006811 | Monoatomic ion transport | 74 | 41.92 | 1.77 | 2.7E-05 |
|  | GO:0006810 | Transport | 198 | 136.81 | 1.45 | 8.07E-07 |
|  | GO:0051234 | Establishment of localization | 199 | 138.31 | 1.44 | 9.09E-07 |
|  | GO:0051179 | Localization | 200 | 138.87 | 1.44 | 9.27E-07 |
|  | GO:0034220 | Monoatomic ion transmembrane transport | 60 | 34.25 | 1.75 | 2.03E-04 |
|  | GO:0055085 | Transmembrane transport | 182 | 120.72 | 1.51 | 2.33E-07 |
|  | GO:0009987 | Cellular process | 594 | 539.20 | 1.10 | 1.77E-04 |
|  | GO:0071702 | Organic substance transport | 133 | 90.58 | 1.47 | 4.74E-05 |
|  | UNCLASSIFIED | Unclassified | 133 | 184.54 | 0.72 | 4.49E-05 |
|  | GO:0006396 | RNA processing | 6 | 22.65 | 0.26 | 1.73E-04 |
|  | GO:0043170 | Macromolecule metabolic process | 136 | 194.27 | 0.70 | 5.15E-06 |
|  | GO:0006399 | tRNA metabolic process | 4 | 18.72 | 0.21 | 1.93E-06 |
|  | GO:0034660 | ncRNA metabolic process | 6 | 26.58 | 0.23 | 9.72E-06 |
|  | GO:0051301 | Cell division | 2 | 15.16 | 0.13 | 1.83E-04 |
| Molecular function | GO:0022857 | Transmembrane transporter activity | 159 | 103.50 | 1.54 | 7.48E-07 |
|  | GO:0005215 | Transporter activity | 159 | 105.00 | 1.51 | 1.49E-06 |
|  | GO:0140640 | Catalytic activity, acting on a nucleic acid | 27 | 56.33 | 0.48 | 4.29E-05 |
| Cellular component | GO:1902495 | Transmembrane transporter complex | 82 | 52.97 | 1.55 | 4.50E-04 |
|  | GO:0098796 | Membrane protein complex | 98 | 63.07 | 1.55 | 1.11E-04 |
|  | GO:0016020 | Membrane | 350 | 281.86 | 1.24 | 7.08E-06 |
|  | GO:0005886 | Plasma membrane | 301 | 239.94 | 1.25 | 2.92E-05 |
|  | GO:0071944 | Cell periphery | 334 | 261.65 | 1.28 | 1.24E-06 |

**Fig. S3.**
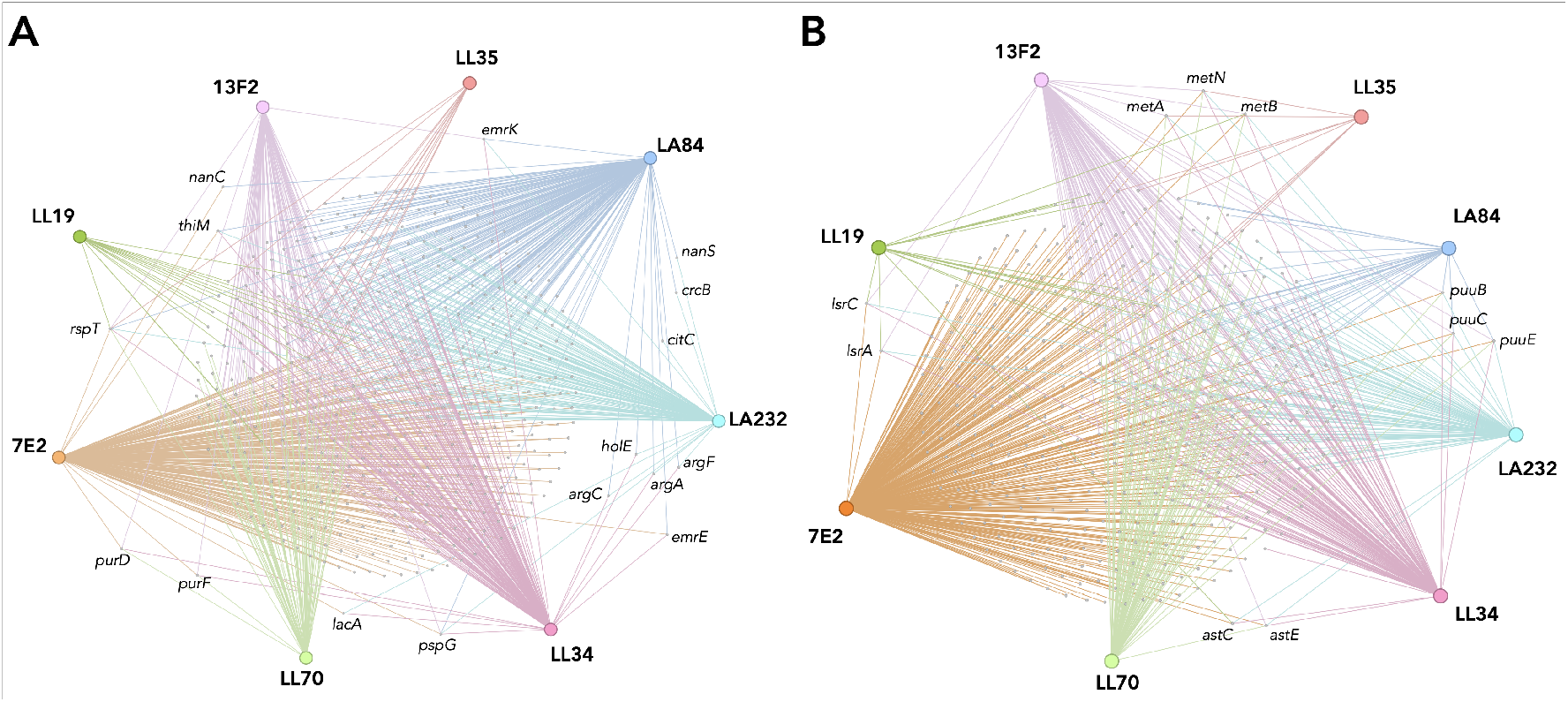
Genes of known function that are **A** up- or **B** downregulated in the eight transconjugants. Nodes depict genes, edges show the transconjugant(s) that they were significantly differentially expressed in. Select genes are highlighted, transconjugants are labelled in bold.

## Notes

### Competing Interest Statement

The authors have declared no competing interest.

